# Neurotox: Deep learning decodes conserved hallmarks of neurotoxicity across venomous species

**DOI:** 10.64898/2026.03.07.710323

**Authors:** Anas Bedraoui, Salim El Mejjad, Salwa Enezari, Fatima Zahra El Hajji, Jacob A. Galan, Rachid El Fatimy, Tariq Daouda

## Abstract

Neurotoxic proteins drive the most pathophysiological effects of animal envenomation, yet it remains unclear whether neurotoxicity is encoded directly within the protein sequence or emerges from higher-order structure binding and interactions with their target receptor. To address this, we developed Neurotox, a sequence-based deep learning framework trained on 200,000 curated protein sequences, with balanced representation of neurotoxic and non-neurotoxic proteins across taxa, achieving high classification accuracy (96%) with strong performance on unseen toxin families. We further introduced a controlled sequence-representation warping strategy that selectively perturbs neurotoxicity-relevant features, inducing a systematic loss of predicted neurotoxicity while preserving primary sequence identity. Structural modeling using AlphaFold 3 showed that, for most top-ranked toxins, warping disrupted β-sheet architectures and reduced interface precision, with all top candidates showing highly significant effects (p < 0.0001). These structural changes were accompanied by recurrent cysteine-centered substitutions, implicating disruption of conserved disulfide frameworks. A single exception retained its global fold (Cα RMSD = 2.8 Å), maintained low PAE, high pLDDT, and high pDockQ scores, and preserved a close arginine-glutamate contact (Arg53-Glu75), yet still exhibited marked attenuation of predicted neurotoxicity. These results suggest that neurotoxicity arises from distributed sequence features that shape secondary-structure organization and receptor interaction, rather than from isolated contact residues alone.

**Teaser:** Deep learning suggests the identification of neurotoxicity hallmarks directly from amino acid sequences across diverse species and toxin families.

## Introduction

Venom-derived neurotoxins represent a fundamental class of bioactive proteins with lasting impact on neurobiology, pharmacology, and clinical medicine (1–7). These molecules disrupt neuromuscular transmission, autonomic signaling, and central nervous system function through highly specific interactions with ion channels and neurotransmitter receptors, leading to paralysis, respiratory failure, cardiac instability, and death in severe envenomation cases (8–10). Neurotoxins have also served as indispensable research tools, supporting detailed investigation of neuronal excitability and synaptic transmission and informing the development of several therapeutics (5,6).

Neurotoxins produced by evolutionarily distant venomous organisms, including snakes, scorpions, spiders, cone snails, and sea anemones, frequently act on the same molecular targets despite substantial sequence divergence (12–16). Although these neurotoxins converge on similar molecular targets, they span diverse protein families and molecular formats, ranging from short peptides to larger proteins with distinct lengths, sizes, and architectures, yet bind specific receptors to produce neurotoxic effects. This combination of shared functional outcome and extreme molecular diversity complicates efforts to identify the sequence features that specify neurotoxicity. Despite extensive biochemical, electrophysiological, and structural study, the sequence-level determinants of neurotoxicity remain poorly defined.

Clinicians still rely primarily on antivenoms that neutralize toxins through antibody recognition, yet these treatments often show limited efficacy against fast-acting neurotoxins, exhibit narrow species specificity, and fail to prevent early neurophysiological damage once toxins bind their targets (17,18). These limitations underscore the need to understand how neurotoxic activity is specified at the molecular level, particularly at toxin-receptor binding interfaces, as this knowledge is critical for rational antivenom design and improving cross-neutralization of structurally related neurotoxins. The field lacks consensus on whether neurotoxicity is governed by discrete sequence elements that directly mediate receptor binding, broader sequence features that shape interaction surfaces, or properties that emerge only after three-dimensional folding.

Extensive experimental work has established that neurotoxicity manifests when venom components engage specific neuronal receptors or ion channels. In snake venoms, neurotoxic effects arise primarily from high-affinity interactions with nicotinic acetylcholine receptors at the neuromuscular junction, leading to rapid paralysis (19). Sea anemone toxins disrupt neuronal signaling through interactions with voltage-gated sodium channels (20), while scorpion and spider neurotoxins modulate sodium, potassium, or calcium channels to alter neuronal excitability (15,21,22). Cone snail venoms act with exceptional specificity through short, disulfide-rich peptides that target ion channels and receptors to produce profound neurophysiological effects (23). Bacterial botulinum neurotoxins induce paralysis through highly specific interactions with presynaptic membrane receptors followed by intracellular cleavage of synaptic proteins that block neurotransmitter release (24). Across these diverse systems, experimental studies have defined neurotoxicity primarily through toxin-receptor interaction and downstream physiological disruption, most often by examining individual toxins or closely related families in isolation (25). This focus has yielded detailed mechanistic insight for specific toxin-receptor pairs but has not established a unifying framework capable of accurately identifying neurotoxic function across structurally and evolutionarily distinct proteins. Thus, it remains unclear whether neurotoxins share common features that predispose them to engage neuronal receptors and cause neurotoxic effects across species, or whether neurotoxicity arises only when individual toxins bind their targets and adopt specific structures.

Recent progress in artificial intelligence has begun to change how researchers analyze protein sequences and structures (5,26–28). Protein language models learn statistical patterns from large collections of amino acid sequences and extract functional information directly from sequence alone (29–31). These models support protein classification and functional prediction without requiring prior structural or experimental data. In parallel, AlphaFold, a deep learning system for protein structure prediction, provides high-accuracy three-dimensional models from sequence, making structural information rapidly accessible (26,32). These advances simplify the selection of candidate proteins for experimental study by prioritizing sequences with predicted functional relevance and providing structural hypotheses before laboratory validation. In venom research, such approaches have supported toxin annotation and structure prediction for proteins lacking experimentally determined structures (33), yet they have rarely been used to test whether neurotoxic activity can be inferred as an intrinsic property of protein sequence rather than as a consequence of receptor binding alone.

We introduce Neurotox, a deep learning framework that learns and interrogates global sequence-level features associated with neurotoxicity. We trained Neurotox on a rigorously curated dataset comprising more than 200,000 protein sequences after preprocessing, with balanced representation of neurotoxic and non-neurotoxic proteins and evaluation on out-of-distribution data, achieving 96% classification accuracy. To test whether the model relied on genuine biological signal rather than dataset correlations, we generated for each neurotoxin a corresponding low-or non-neurotoxic variant through controlled perturbation or warping, which selectively modifies sequence to reduce the probability of the protein being classified as neurotoxic. From these results, we selected top-ranked neurotoxins for structural analysis based on three criteria: high neurotoxicity probability in the original model, strong reduction of predicted neurotoxicity after warping, and high sequence similarity between native and warped variants. We then combined these sequence perturbations with three-dimensional structure prediction and neurotoxin-receptor interaction modeling using AlphaFold 3 to examine how loss of predicted neurotoxicity propagated to molecular interfaces. Our analysis shows that reduced neurotoxic signal coincides with diminished receptor-binding precision while global protein folds remain intact. These findings support the existence of shared sequence-encoded determinants of neurotoxic activity and provide a framework for improving neurotoxin annotation and guiding safer neuroactive drug and antivenom design.

## Results

### Embedding Warping Selectively Disrupts Neurotoxic Hallmarks While Preserving Global Structural and Phylogenetic Organization

Having identified neurotoxins that retain high structural similarity while exhibiting a marked loss of neurotoxic signal after embedding warping, we next examined how this disruption manifests at the level of individual proteins **(Fig. 1)**. Five top-ranked neurotoxins, spanning distinct venomous taxa, were selected based on the criteria mentioned above. Comparison of neurotoxicity probabilities before and after warping revealed a consistent and substantial reduction across all five proteins, with original sequences exhibiting high neurotoxicity scores exceeding 90% **(Fig. 1A)** and their warped counterparts showing markedly reduced probabilities below 60% **(Fig. 1B)**, despite only minor structural changes and preservation of the overall three-dimensional fold, with the exception of P80494 (muscarinic toxin alpha) from elapid snake venom, which retained its native structure.

**Fig. 1.**
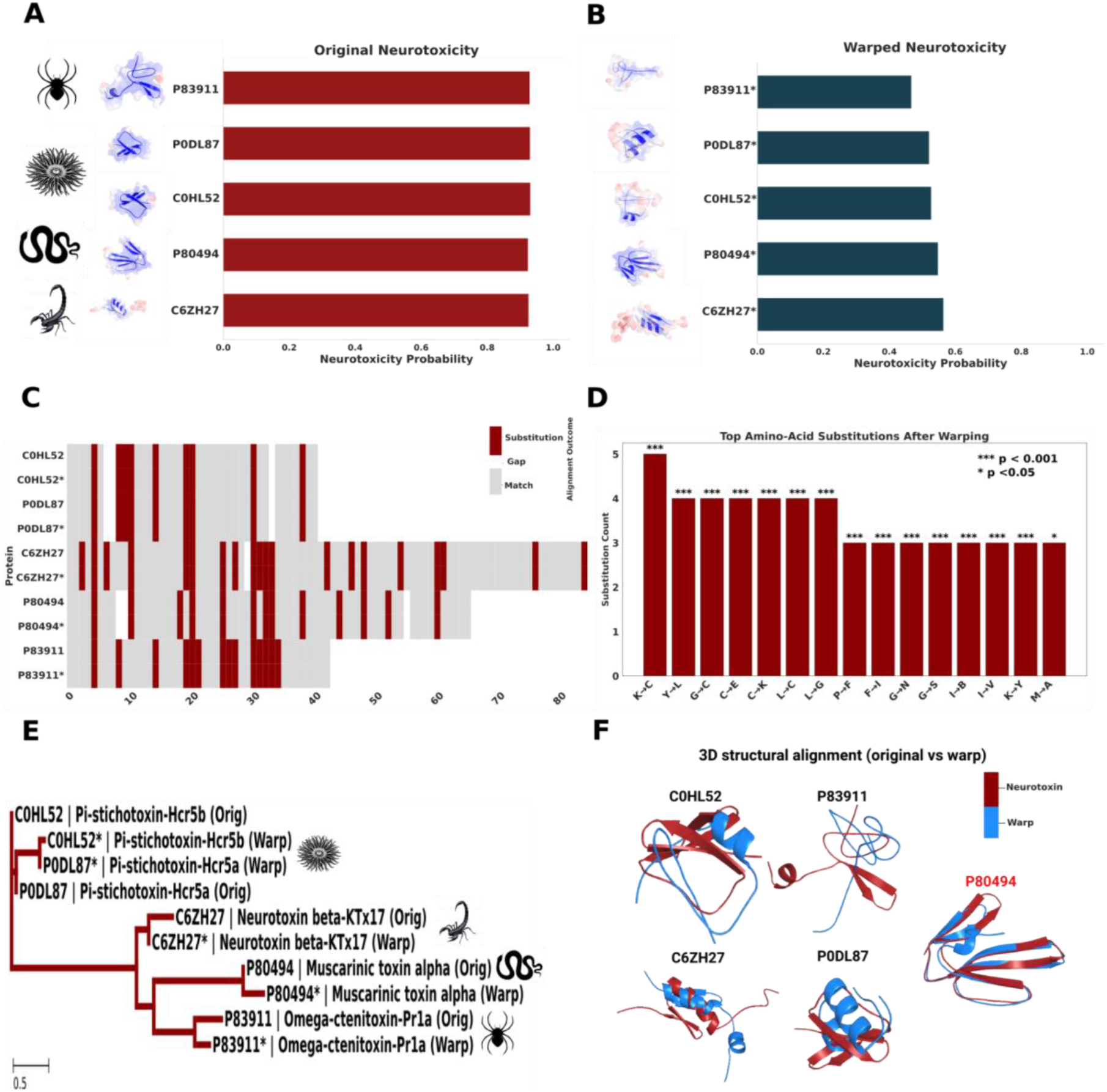
Functional, Structural, and Topological Effects of Sequence Warping in Top-Ranked Neurotoxin Proteins. (A) Predicted neurotoxicity probabilities for the five selected venom proteins prior to warping, showing uniformly high confidence scores across all sequences. (B) Corresponding neurotoxicity probabilities after sequence warping, revealing a consistent and substantial reduction in predicted neurotoxic activity (all <60%) while preserving high sequence similarity (>75%). (C) Local pairwise alignment heatmap comparing original and warped sequences. Conserved aligned positions are shown in grey, whereas amino acid substitutions introduced by warping are highlighted in red, illustrating the distribution of sequence perturbations along each protein. (D) Frequency of amino acid substitutions observed after sequence warping, reported directionally from the original residue to the substituted residue. Enriched substitution types highlight recurrent, statistically significant residue changes associated with loss of neurotoxicity, with significance levels indicated (* p < 0.05, *** p < 0.001). (E) Topological relationships among original and warped protein sequences demonstrate that each warped variant clusters as a sister to its corresponding original, indicating preservation of lineage-level sequence similarity despite marked functional attenuation. (F) Three-dimensional structural alignments of original (red) and warped (blue) proteins illustrating β-sheet disruption and protein-specific secondary-structure reorganization induced by sequence warping.

To localize where neurotoxic hallmarks are disrupted within individual sequences, we next examined local pairwise alignments between original and warped versions of each selected neurotoxin **(Fig. 1C)**. Although overall sequence similarity was preserved, substitutions were not randomly distributed but showed convergent patterns across proteins. Despite differences in sequence length, multiple toxins exhibited substitutions concentrated within corresponding N-terminal to early central segments of the alignment, indicating common regions of perturbation across otherwise distinct sequences. To further examine how embedding warping perturbs neurotoxic function at the residue level, directional amino acid substitution enrichment was assessed within a high-confidence loss-of-neurotoxicity subset (sequence similarity ≥75%, original probability ≥0.90, warped probability ≤0.60). Substitutions were reported directionally from the original to the warped residue and evaluated using two-sided Fisher’s exact tests, as all substitution counts were sparse (n < 5) **(Fig. 1D)**. This shows that the substitution patterns observed in the top-ranked neurotoxins recapitulate the dominant perturbation modes identified at the global scale. Importantly, the overlap between these two substitution spectra is non-random and structurally informative.

Topological relationships between original and warped toxin sequences were inferred from multiple sequence alignment using a maximum likelihood framework implemented in IQ-TREE under the best-fit substitution model selected by ModelFinder (PMB+G4). The resulting rectangular phylogram **(Fig. 1E)** shows that each warped sequence clusters as a well-supported sister to its corresponding original toxin, indicating that sequence warping preserves lineage-level evolutionary structure. No warped sequences formed independent cross-family clusters; instead, all sequences remained organized by toxin family and biological origin. Although branch lengths reflect measurable warp-induced divergence, this divergence remains substantially smaller than inter-family separation, demonstrating that sequence warping perturbs functional determinants while preserving global scaffold identity and family-level organization.

Three-dimensional structural alignment between AlphaFold3-predicted models of original and warped sequences was quantified using Cα root-mean-square deviation (RMSD) following pairwise three-dimensional structural alignment **(Fig. 1F)**. All selected neurotoxins share a common β-sheet–based structural scaffold in their native state (red). Following sequence warping (blue), this β-sheet framework was disrupted in most toxins, often accompanied by a transition toward α-helical or loop-dominated conformations. Specifically, C6ZH27, P0DL87, and C0HL52 exhibited loss of β-sheet elements with emergence of α-helical segments and loop elongation, while P83911 showed β-sheet destabilization and collapse into a flexible, loop-rich structure. In contrast, P80494 retained all three β-sheets with minimal backbone deviation (Cα RMSD = 2.8 Å), indicating exceptional structural robustness despite sequence perturbation.

### AlphaFold 3 Prediction of Neurotoxin-Receptor Interface Stability

AlphaFold 3 was used to predict three-dimensional structures of top ranked neurotoxins, their corresponding neuroreceptors, and their multimeric complexes in order to examine how embedding warping alters neurotoxin-receptor binding at the structural level. For each toxin-receptor pair, five independent AlphaFold 3 models were generated, and both global and interface-level confidence metrics were calculated to quantify structural reliability, interface precision, and interaction consistency **(Fig. 2)**.

**Fig. 2.**
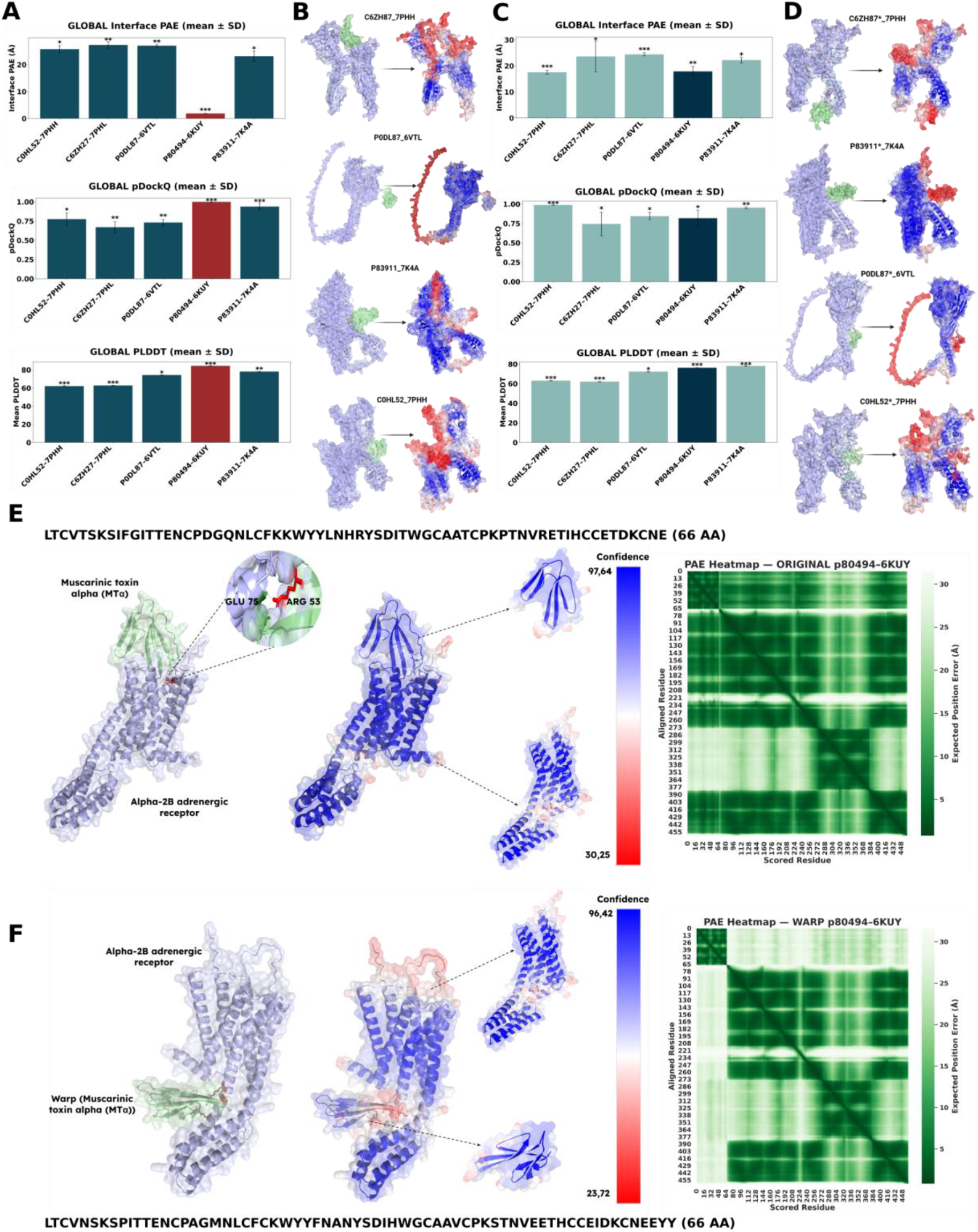
Structural Reliability of Top-Ranked Neurotoxin-Receptor Interactions Assessed by AlphaFold 3. (A) Global interface confidence metrics for native neurotoxin-receptor complexes derived from five independent AlphaFold 3 models per interaction. Interface predicted aligned error (PAE), pDockQ, and mean pLDDT are reported as mean ± SD. The interaction highlighted in red is uniquely distinguished by exceptionally low interface PAE and high pDockQ and pLDDT values, identifying it as the most structurally precise and reliable complex. (B) Structural visualization of native neurotoxin–receptor complexes exhibiting elevated interface PAE. Toxins are shown in green and receptors in light blue, with per-residue confidence mapped using a blue-white-red scale, where red indicates low local confidence. In all four complexes, low-confidence (red) regions dominate the predicted binding interfaces, reflecting weakly constrained extracellular interaction geometries. (C) Global interface confidence metrics for warp-engineered toxin-receptor complexes. Interface PAE values increase substantially across all interactions, whereas pDockQ and pLDDT remain relatively preserved, indicating selective degradation of interface precision without global unfolding. (D) Structural visualization of warped toxin–receptor complexes. Warped toxins adopt shifted and heterogeneous orientations on the receptor surface, with extensive, red-colored regions concentrated at the predicted interfaces, indicating increased positional uncertainty and diffuse binding modes. (E) High-resolution PyMOL visualization of the top-ranked native P80494-6KUY interaction reveals a compact and rigid extracellular binding mode stabilized by a short-distance electrostatic contact. Uniformly high local pLDDT values and a pronounced low-PAE valley indicate exceptional interface precision. (F) Corresponding visualization of the warped P80494-6KUY interaction shows loss of the canonical binding pose, with multiple contacts occurring in low-confidence (red) regions characterized by reduced pLDDT and elevated PAE, indicating destabilized and poorly constrained receptor interaction.

Analysis of the original toxin-receptor complexes revealed pronounced heterogeneity in interface quality across interactions **(Fig. 2A)**. All four complexes exhibited elevated interface predicted aligned error (PAE), exceeding 20 Å, indicative of flexible, weakly constrained extracellular binding geometries. In contrast, one interaction was uniquely distinguished by a low interface PAE, reflecting a highly precise and spatially constrained toxin-receptor interface between muscarinic toxin alpha (P80494) and its corresponding neuroreceptor, the α2A adrenergic receptor (6KUY). This same interaction also displayed the highest pDockQ and global predicted local distance difference test (pLDDT) values, indicating both strong docking confidence and highly stable folding of the toxin and receptor. The convergence of low interface PAE with high pDockQ and pLDDT identifies this interaction as the most structurally reliable neurotoxin-receptor complex in the top ranked candidates.

Structural visualization of the remaining original complexes provided direct insight into the origin of their elevated interface uncertainty **(Fig. 2B)**. In these four complexes, the predicted binding sites were dominated by regions colored red in the confidence mapping, corresponding to residues with low local confidence and high positional uncertainty. Notably, these red regions were concentrated precisely at the toxin-receptor interface rather than being distributed across the protein surface, except for the P0DL87-6VTL pair, for which the predicted complex appeared to be hallucinatory. Such behavior reflects a known limitation of AlphaFold-based multimer predictions, particularly for weak, transient, or non-cognate interactions, where insufficient co-evolutionary or structural constraints can lead the network to impose an artificial or hallucinatory interface (34). The toxins adopted variable orientations relative to the receptor extracellular domains, and interface contacts were dispersed across broad, flexible regions instead of forming a compact, well-defined binding site. This pervasive low-confidence (red) signal at the binding interface explains the high interface PAE values observed for these complexes and indicates that, despite reasonable global folding, the precise geometry of toxin engagement is poorly resolved.

We next assessed warp toxin variants using the same pipeline described above. Across all complexes, interface PAE values increased substantially relative to their original counterparts, exceeding 18 Å **(Fig. 2C)**. This increase reflects a systematic loss of interface precision following embedding perturbation. Importantly, pDockQ values remained moderate to high in several warped complexes, indicating that docking confidence metrics alone can overestimate interaction reliability when interface geometry is poorly constrained. Mean pLDDT values remained largely unchanged, confirming that embedding warping does not induce global unfolding of either the toxin or the receptor. Structural inspection of the warped complexes further reinforced this interpretation **(Fig. 2D)**. In contrast to the original toxins, reduced model confidence was predominantly localized to the warped toxin and concentrated at the toxin-receptor contact interface. Warped toxins adopted shifted and heterogeneous orientations on the receptor surface, with interface regions dominated by red-colored residues at the binding site, indicating low local structural confidence.

To examine these effects at atomic resolution, we focused only on the top-ranked original interaction (P80494-6KUY). PyMOL visualization of the highest-confidence AlphaFold 3 model revealed a compact and rigid extracellular binding mode **(Fig. 2E)**. In stark contrast to the four weaker complexes, the binding interface was largely devoid of red regions and instead dominated by uniformly high-confidence residues (blue). The interaction was stabilized by a single short-distance electrostatic contact between ARG53 of muscarinic toxin α (P80494) and GLU75 of the α2A-adrenergic receptor (6KUY) at 2.38 Å. This minimal but highly specific contact network coincided with uniformly high local pLDDT values, reaching a maximum of 97.53, and a pronounced low-PAE valley in the confidence heatmap. These indicate exceptional positional certainty and a highly optimized toxin-receptor interface.

In contrast, PyMOL inspection of the best-ranked warped model of the same toxin-receptor pair revealed a marked collapse of interface precision **(Fig. 2F)**. The engineered toxin no longer occupied the same exact canonical extracellular binding pose and instead adopted a shifted and more flexible orientation on the receptor surface. Multiple short-distance contacts were observed, but these were clustered within regions colored red, reflecting low local confidence and elevated PAE. The emergence of extensive red patches at the interface indicates substantial conformational uncertainty and weakened binding precision, despite preservation of the overall fold **(Fig. 1E)**. This transition from a compact, high-confidence interface to a diffuse, low-confidence interaction surface underscores the selective disruption of functional binding geometry induced by embedding warping. This selective degradation of interface geometry provides a structural explanation for the loss of predicted neurotoxicity observed following embedding warping **(Fig. 1B)**.

## Discussion

Neurotoxicity arises from highly specific interactions between venom components and molecular targets in the nervous system (25). These interactions depend on a balance between conserved structural scaffolds and finely tuned surface residues that together define receptor recognition and binding affinity (35). Once bound, neurotoxins disrupt normal neuronal signaling by altering ion channel conductance, blocking neurotransmitter receptors, or interfering with synaptic vesicle release (19). These molecular perturbations propagate through neuronal circuits by modifying the activity of voltage-gated ion channels that regulate neuronal excitability, including sodium (Na_v_) channels controlling membrane depolarization, potassium (K_v_) channels governing repolarization, and calcium (Ca_v_) channels that trigger neurotransmitter release at synaptic terminals, thereby altering neuromuscular transmission and neuronal signaling (36–39).

Many venom neurotoxins exhibit remarkable receptor specificity. For example, snake three-finger toxins (3FTx) bind nicotinic acetylcholine receptors (nAChRs) at the neuromuscular junction with nanomolar affinity, preventing acetylcholine binding and producing rapid postsynaptic paralysis (40). Other venom systems target ion channels responsible for neuronal excitability. Cone snail conotoxins are highly selective ligands for voltage-gated sodium, potassium, and calcium channels, as well as N-methyl-D-aspartate receptor (NMDA) and nicotinic receptors, making them among the most pharmacologically diverse neuropeptide families known (41–43). Sea anemone toxins similarly modulate voltage-gated potassium and sodium channels, often altering channel gating kinetics and action potential propagation (44,45).

Arthropod venoms provide additional examples of highly specialized neurotoxins. Scorpion toxins are well known for targeting voltage-gated sodium (Na_v_) and potassium (K_v_) channels, where α- and β-toxins alter channel activation or inactivation states, leading to prolonged depolarization and neuronal hyperexcitability (46,47). Spider venoms contain numerous inhibitor cystine knot (ICK) peptides that modulate ion channels involved in nociception and synaptic transmission, including voltage-gated calcium channels and mechanosensitive channels (48). Many spider toxins act as gating modifiers that stabilize specific channel conformations, thereby altering the probability of channel opening (49,50).

At the extreme end of neurotoxic potency, bacterial toxins such as botulinum neurotoxins act through a distinct mechanism by enzymatically cleaving SNARE proteins required for synaptic vesicle fusion (51). This process prevents acetylcholine release at presynaptic terminals, producing flaccid paralysis. Despite mechanistic differences across taxa, these systems converge on a common functional outcome: disruption of neuronal communication.

Venoms are therefore complex biochemical mixtures in which multiple toxins can act simultaneously on related physiological targets. Synergistic interactions among toxin families frequently amplify toxicity by affecting several stages of neuromuscular signaling. For instance, presynaptic phospholipases A_2_ may facilitate toxin diffusion and membrane disruption, while postsynaptic α-neurotoxins block receptor activation (52,53). Such combinatorial strategies allow venom systems to rapidly incapacitate prey through coordinated interference with synaptic transmission.

Biochemically, snake venoms consist predominantly of proteins and peptides, typically representing approximately 90-95% of the dry venom mass, although this proportion varies among taxa and venom systems (54). These molecules range from short disulfide-rich peptides to larger multidomain enzymes that may form dimers or higher-order complexes capable of targeting specific receptors. Despite their diversity, many toxins share conserved structural scaffolds such as the three-finger fold, inhibitor cystine knot (ICK), Kunitz domains, and phospholipase structures that stabilize their conformation and enable precise receptor binding (52,55–57).

This combination of shared physiological outcome and extensive molecular diversity complicates efforts to identify the sequence features that define neurotoxicity. While structural motifs provide clues to function, subtle variations in surface residues can dramatically alter receptor specificity and pharmacological activity. Consequently, despite decades of biochemical, electrophysiological, and structural studies, the sequence-level determinants governing neurotoxic activity remain incompletely understood. These challenges have motivated the development of computational approaches designed to infer toxin function directly from sequence data and large curated venom datasets.

Several computational tools have addressed toxin and neurotoxicity prediction using diverse methodological strategies. ToxTeller applies logistic regression, SVM, random forest, and XGBoost to curated peptide datasets, achieving 85.5% accuracy, 0.930 AUC, and 0.721 MCC on an independent ≤40% similarity test set, though performance remains feature-dependent (58). NTXpred, trained on 582 experimentally validated neurotoxins, reported up to 97.72% accuracy using SVM under five-fold cross-validation, with source and functional classification reaching 92– 95% when incorporating PSI-BLAST profiles, however, it was developed on a relatively small dataset and lacked independent family-level validation (59). CAPTP combines convolutional layers, multi-head self-attention, and contrastive learning, achieving 91.59% balanced accuracy and 0.811 MCC on independent peptide datasets (60). NeuTox 2.0 targets small-molecule neurotoxicity via multimodal molecular feature fusion, reporting AUC 0.97 across chemical toxicity endpoints (60). CSM-Toxin leverages ProteinBERT-based language embeddings from primary sequences, attaining MCC up to 0.66 on cross-validation and blind tests (61). ToxPre-2L uses multi-view tensor learning with low-rank constraints for peptide toxicity and subtype prediction, achieving 0.86 accuracy under 10-fold cross-validation (62).

Relative to prior tools, Neurotox advances neurotoxicity prediction in scale, evaluation rigor, and structural interrogation. Unlike NTXpred (n = 582) or peptide-focused models such as ToxTeller, CAPTP, and ToxPre-2L, Neurotox was trained on 200,000 curated protein sequences with balanced neurotoxic and non-neurotoxic representation across taxa. Importantly, 20% of toxin families were completely excluded during training, enabling strict family-level validation, and yielding 96% accuracy on unseen families, a more stringent strategy than similarity-based splits or cross-validation alone. Beyond sequence-level classification, Neurotox extends into three-dimensional structural space. Using AlphaFold 3, we demonstrated that controlled sequence-representation warping disrupted β-sheet architectures and reduced interface precision with highly significant effects (p < 0.0001), frequently associated with cysteine-centered substitutions affecting conserved disulfide frameworks (Fig. 1 D, F). Even in a structurally preserved outlier (Cα RMSD = 2.8 Å, high pLDDT and pDockQ), predicted neurotoxicity markedly decreased (Fig. 1B). In contrast, most existing tools remain confined to sequence-or descriptor-level prediction without systematic 3D structural validation. Collectively, Neurotox integrates large-scale family-aware training with structural modeling, providing both predictive performance and mechanistic insight into sequence-encoded neurotoxicity.

Building on this integrative framework, our findings support a broader conceptual interpretation of neurotoxic function (63). Neurotoxins from distantly related species can converge on similar physiological effects because they share common representational and structural principles, even when their primary sequences and evolutionary origins differ (57,64). Neurotox captures these principles at the sequence-representation level, and controlled perturbation of these representations disrupts the neurotoxic signal without requiring wholesale loss of sequence identity. This shifts attention away from isolated residues or short linear motifs and toward higher-order organization of distributed sequence features that collectively govern functional specificity.

Residue-level patterns emerging from representation perturbation suggest that certain amino acid classes play recurring roles in neurotoxic activity (Fig. 1D). Cysteines, through their role in disulfide bond formation, anchor structural frameworks that constrain loop geometry and surface presentation. Glycine residues contribute flexibility that supports precise conformational sampling, while charged and aromatic residues shape electrostatic and hydrophobic interactions at receptor-facing surfaces. Neurotoxicity therefore appears to arise from the coordinated placement of these residues rather than their presence in isolation, offering a unifying explanation for functional conservation across venom families. Structural modeling extends this interpretation by highlighting the importance of interaction precision rather than binding presence alone. Native neurotoxins tend to form compact, well-defined interfaces characterized by low positional uncertainty, consistent with high specificity and potency.

Among the top-ranked toxins analyzed, muscarinic toxin alpha (P80494) from *Dendroaspis polylepis polylepis* represents a notable structural outlier (Fig. 2). This 66-amino acid, disulfide-rich protein is experimentally validated at the protein level and is known to bind selectively and with high affinity to the ADRA2B receptor subtype, with additional reported nanomolar affinities toward multiple muscarinic receptor subtypes (65–68). In the present study, structural complex modeling was performed using the closely related α2A adrenergic receptor (ADRA2A), for which high-resolution structural templates are available (69). Although direct experimental validation of P80494 binding to ADRA2A remains limited, the strong structural homology between α2A and α2B subtypes provides a rational framework for exploring subtype-level interaction conservation (70). Despite controlled representation warping, the toxin retained remarkable structural stability. Structural alignment between the original and warped models yielded a low Cα RMSD of 2.38 Å, indicating minimal global conformational deviation. AlphaFold confidence metrics remained high, with pLDDT values reaching 97.53, supporting preservation of the canonical disulfide-constrained fold. Remarkably, however, its predicted neurotoxicity probability dropped substantially following representation perturbation. This apparent decoupling between global structural conservation and predicted functional activity suggests that subtle sequence perturbations may disrupt critical structural determinants underlying neurotoxic function.

These findings support the conclusion that neurotoxicity hinges not just on primary sequence identity or global fold preservation, but on sequence-driven perturbations that reshape secondary structure organization, ultimately modulating neural receptor interactions. This principle is echoed in the literature: for instance, snake venom neurotoxins like erabutoxin b, α-cobratoxin, and α-bungarotoxin maintain β-sheet cores stabilized by specific disulfides, where guanidinium hydrochloride titrations reveal sequential unfolding starting at peripheral strands but preserving the Trp-embedded center longest, directly linking local secondary disruptions to functional loss (71). Similarly, amyloid-β oligomers show toxicity scaling with β-sheet content (tetramers far exceeding monomers), while botulinum neurotoxin/A1 mutants retain overall folds yet exhibit up to 70,000-fold potency reductions from catalytic domain remodeling (72–74). These examples highlight secondary structure plasticity as a tunable switch for neurotoxicity, offering promising avenues for scaffold-targeted therapeutic interventions.

From this perspective, neurotoxins from distantly related species can converge on similar physiological effects because they share common representational and structural principles, even when their sequences and evolutionary origins differ (75). Neurotox captures these principles at the sequence-representation level, and controlled perturbation of these representations disrupts neurotoxic signal without requiring wholesale loss of sequence identity. This shifts attention away from single residues or motifs and toward higher-order organization of sequence features that govern functional specificity.

Despite these insights, several limitations should be considered when interpreting these results. Representation warping is an abstract computational operation and does not correspond directly to physical mutations. Experimental validation will therefore be required to determine whether the substitutions highlighted here reproduce loss of neurotoxicity in vitro or in vivo. Structural predictions rely on modeled confidence metrics rather than direct measurements of receptor binding affinity or physiological activity, and receptor selection may not capture the full diversity of toxin targets. In addition, currently available neurotoxin datasets remain relatively limited and unevenly distributed across venomous taxa, which may constrain model generalization and bias predictions toward well-characterized toxin families. To partially mitigate this imbalance, we implemented a family-wise oversampling strategy to approximate equal representation across toxin families, as described in the Methods. While this approach helps reduce phylogenetic and functional skew in the training data, it does not fully compensate for the limited diversity of experimentally characterized neurotoxins, and broader taxonomic sampling will be necessary to better capture the full landscape of neurotoxic proteins.

## Materials and Methods

### Sequence Alignment and Selection of High-Confidence Warped Hits

To characterize how embedding warping altered sequence-level features while preserving primary sequence identity, we performed a systematic alignment and variability analysis comparing original sequences with their warped counterparts. All analyses were conducted at the individual-protein level and restricted to proteins for which both original and warped representations were available.

Warped sequence variants generated across warped model instances were grouped by UniProt identifier, collapsing isoform suffixes when present. For each protein, the original amino acid sequence was used as a reference, and all associated warped sequences were compared against it. Sequence identity was quantified using position-wise comparisons after padding sequences to equal length, yielding a percent identity score for each warped variant. For downstream analyses, the warped sequence with the highest identity to the original was selected as the representative variant, ensuring maximal preservation of primary sequence similarity. Minor terminal artifacts introduced during warping were corrected by restoring original residues at sequence termini when necessary.

To assess positional variability induced by warping, all warped sequences for a given protein were aligned by position to the original sequence. For each residue position, the frequency of amino acid substitutions across warped variants was computed, and positions at which the most frequent warped residue differed from the original were classified as variable sites. Substitution frequency and percentage were recorded for each variable position, yielding a per-protein positional variability profile. In parallel, local pairwise alignments between original and warped sequences were performed using a substitution-matrix-based alignment approach to explicitly extract residue-level substitution events while accounting for local insertions and deletions. These alignments enabled direct annotation of amino acid changes introduced by warping, and directional enrichment of specific amino acid substitutions was evaluated using contingency-table analysis, applying Fisher’s exact test for low-count events and χ^2^ tests for higher-frequency substitutions to determine statistically significant deviations from background frequencies.

Sequence stability was summarized at the protein level by aggregating positional variability statistics, including the number of variable positions, mean variability, maximum variability, and an overall stability score defined as the complement of mean positional variability. Substitution events were further aggregated across all proteins to generate a global substitution spectrum, highlighting recurrent original-to-warped amino acid exchanges, and enabling identification of dominant physicochemical trends disrupted by warping. To identify top candidates for structural and functional follow-up, proteins were filtered using a combined sequence- and prediction-based criterion. Proteins were retained if they satisfied three conditions: high sequence similarity between original and warped sequences (>75% identity), high neurotoxicity probability under the original model (>90%), and a substantial reduction in neurotoxicity probability under the warped model (<60%). This strategy isolated proteins that ranked highly in the original model but dropped sharply in the warped model, while maintaining high sequence similarity and identity, thereby enabling focused downstream structural and mechanistic analyses.

### Structural Modeling and Interface Evaluation

To determine whether embedding warping alters neurotoxin–receptor interactions at the structural level, three-dimensional structure prediction and quantitative complex evaluation were performed for top-ranked protein hits identified in the sequence-level analyses. All procedures were applied in parallel to original and warped sequences, using identical receptors, prediction parameters, and evaluation criteria to enable direct comparison.

For each selected protein, mature toxin sequences were manually curated prior to structure prediction. Signal peptides and propeptide regions were removed based on published biochemical annotations and primary literature specific to each toxin family. Only reviewed proteins with well-established sequence boundaries were retained, ensuring that all modeled sequences corresponded to biologically active toxin forms. Curated toxin sequences were submitted to AlphaFold 3 for protein-receptor complex prediction. Receptors were selected based on experimental and structural evidence reported in the literature, and the same receptor was used for both original and warped variants of each toxin. For each toxin-receptor pair, AlphaFold 3 generated five independent complex models, along with associated metadata files reporting predicted aligned error (PAE), inter-chain predicted TM scores (ipTM), and per-residue confidence metrics. All five models were retained for downstream analysis.

Structural confidence and interface quality were quantified using three complementary metrics extracted independently from each AlphaFold 3 model. Interface predicted aligned error (PAE) was obtained from the chain-pair minimum PAE matrices in the AlphaFold 3 output. For each complex, the receptor chain exhibiting the strongest predicted interaction with the toxin chain was identified based on inter-chain ipTM values, and the corresponding interface PAE was extracted. Docking confidence was quantified using pDockQ, computed from inter-chain ipTM values via a logistic transformation. Global structural reliability was assessed using mean per-residue PLDDT, extracted from the B-factor field of the predicted mmCIF structures.

For each toxin-receptor pair, interface PAE, pDockQ, and PLDDT values were aggregated across the five predicted models, yielding mean and standard deviation estimates. Aggregation was performed separately for original and warped complexes. To contextualize individual complexes relative to the full dataset, each complex was statistically compared against all other complexes using Welch’s two-sample t-tests, performed independently for each metric. Resulting p-values were used to identify structurally distinct interactions and were visualized using star-based significance annotations.

All three-dimensional structural visualizations were generated using PyMOL (Schrödinger). Identical visualization protocols were applied to original and warped complexes to ensure direct comparability. Structures were rendered using combined cartoon and surface representations, with toxin chains consistently designated as chain A and receptors as chain B. Toxin chains were colored pale green and receptor chains light blue. Per-residue confidence was visualized by mapping PLDDT values to a red-white-blue color spectrum, where red indicates low confidence, white intermediate confidence, and blue high confidence. This confidence mapping was applied uniformly across all structures. Protein-receptor interfaces were annotated by selecting residues less than 2.5 Å distance cutoff between toxin and receptor chains. Interface residues were displayed in stick representation and color-coded by chain, and atom-atom distance measurements were rendered directly in the structure view for detailed inspection of interaction geometry.

## Notes

### Competing Interest Statement

The authors have declared no competing interest.

